# A template-matching algorithm for laminar identification of cortical recording sites from evoked response potentials

**DOI:** 10.1101/749069

**Authors:** Giulio Matteucci, Margherita Riggi, Davide Zoccolan

**Author notes:** Corresponding author Davide Zoccolan International School for Advanced Studies (SISSA) Via Bonomea, 265 34136 Trieste (TS) ITALY. these authors contributed equally to this work (Giulio Matteucci, Margherita Riggi).

## Abstract

In recent years, the advent of the so-called silicon probes has made it possible to homogeneously sample spikes and local field potentials (LFPs) from a regular grid of cortical recording sites. In principle, this allows inferring the laminar location of the sites based on the spatiotemporal pattern of LFPs recorded along the probe, as in the well-known current source-density (CSD) analysis. This approach, however, has several limitations, since it relies on visual identification of landmark features (i.e., current sinks and sources) by human operators – features that can be absent from the CSD pattern if the probe does not span the whole cortical thickness, thus making manual labelling harder. Furthermore, as any manual annotation procedure, the typical CSD-based workflow for laminar identification of recording sites is affected by subjective judgment undermining the consistency and reproducibility of results. To overcome these limitations, we developed an alternative approach, based on finding the optimal match between the LFPs recorded along a probe in a given experiment and a template LFP profile that was computed using 18 recording sessions, in which the depth of the recording sites had been recovered through histology. We show that this method can achieve an accuracy of 79 µm in recovering the cortical depth of recording sites and a 76% accuracy in inferring their laminar location. As such, our approach provides an alternative to CSD that, being fully automated, is less prone to the idiosyncrasies of subjective judgment and works reliably also for recordings spanning a limited cortical stretch.

**New and noteworthy:** Knowing the depth and laminar location of the microelectrodes used to record neuronal activity from the cerebral cortex is crucial to properly interpret the recorded patterns of neuronal responses. Here we present an innovative approach that allows inferring such properties with high accuracy and in an automated way (i.e., without the need of visual inspection and manual annotation) from the evoked response potentials (ERPs) elicited by sensory (e.g., visual) stimuli.

## Introduction

Most neuronal circuits in the mammalian brain are characterized by a complex spatial organization that is tightly intertwined with their function. In particular, in the cortex, laminar structure is closely linked to the flow of information among different neuronal populations (Adesnik and Naka 2018; Harris and Mrsic-Flogel 2013). Therefore, to fully understand the principles of operation of cortical circuits, it is essential to analyze the activity of single neurons in their spatial context.

Historically, extracellular recordings have been the workhorse method for studying cortical functions and they are still largely used in system neuroscience experiments, although neurophysiologists are increasingly replacing the traditional single electrode approach with multielectrode arrays. The latter allow isolating the waveforms of the action potentials (a.k.a. spikes) fired by many single neurons at once with sub-millisecond temporal precision (Harris et al. 2016) – with the number of recorded units ranging from a few tens to many hundreds, depending on the shape, geometry, technology and materials used to build the array (Chung et al. 2019; Jun et al. 2017). However, such powerful experimental approach has a main limitation: its spatial “blindness”. In fact, looking at the spikes detected on a given channel of an extracellular electrode gives no direct information about the spatial (e.g., laminar) location of the source neurons. Thus, it is not surprising that the challenge of coordinating multielectrode recordings with anatomical information has been recognized in the literature as a fundamental one (Buzsáki et al. 2012; Li et al. 2015). In particular, the most basic anatomical metadata needed to fully make sense of extracellular cortical recordings is the laminar identity of the recorded single units (i.e., the cortical layers in which the recorded neurons sit).

This problem has been partially addressed by the advent of the so-called silicon (or laminar) probes – fork-shaped silicon substrates with several shanks, along which multiple recording sites are placed with a regular spacing (Blanche et al. 2005; Steinmetz et al. 2018). These arrays allow spanning homogeneously the whole cortical thickness or a part of it, thus recording simultaneously spiking signals and “local field potentials” (LFPs) from neurons located in multiple cortical layers. This makes it possible, in principle, to infer the laminar location of the recording sites based on the spatio-temporal pattern of LFPs recorded along the probe, without resorting to laborious and time-consuming histological procedures.

Throughout the years, a technique known as “current source-density” (CSD) analysis has been widely exploited to fulfill this goal in different animal models, from monkeys (Hansen et al. 2011; Schroeder et al. 1998, 2001; Takeuchi et al. 2011) to rodents (Dadarlat and Stryker 2017; Deliano et al. 2018; Happel et al. 2010; Headley and Weinberger 2015; Hoy and Niell 2015; Li et al. 2018; Niell and Stryker 2008; Sieben et al. 2013). CSD exploits the LFP gradients recorded across geometrically-arranged electrode arrays to estimate the extracellular current flow in the surrounding tissue (Nicholson and Freeman 1975). This is done by computing the second spatial derivative of the LFPs recorded along the axis perpendicular to the laminar structure of the cortex. In fact, this quantity is directly proportional to the extracellular current, if one assumes the tissue as being composed of a stack of two-dimensional isopotential planes. The resulting pattern of current sinks and sources across the cortical depth, mainly linked to local synaptic activity (Buzsáki et al. 2012), can thus be used to infer the laminar location of the recording sites by visual inspection. More specifically, a key landmark used in this process is the prominent current sink associated with thalamic afferent inputs impinging into layer 4 (L4) of primary sensory cortices. Such characteristic CSD feature is usually associated with an inversion of the polarity of the stimulus-evoked LFP waveforms – i.e., the evoked response potentials (ERP), a.k.a. visually evoked potentials (VEP) in the case of visual stimulation.

Despite its widespread use, laminar identification through CSD has several drawbacks and limitations. First, the identification of the layers is usually carried out through visual inspection of the CSD pattern. Such reliance on the subjective judgment of the investigator implies a lack of objectivity and standardization, the unknown precision of the inference, and the slowness and laboriousness of the process. Second, although the CSD is meant to enhance the spatial resolution in the localization of the signal’s source, as compared to raw LFPs, this comes at the expense of being more sensitive to imperfections of the electrode array (e.g., unwanted variations of channels impedance along the electrode array that may occur with re-use of the probe or due to fabrication defects). Third, and more importantly, to infer with reasonable confidence the laminar position of the recording sites, it is essential to span a large fraction of the cortical depth, so as to observe the landmark sink in L4. This can be a serious limitation, when silicon probes with tightly packed sites (e.g., 25 µm inter-site spacing) are inserted into the cortex (often with a tilt), so as to densely sample neuronal populations from a specific supragranular or infragranular layer, given that the L4 sink will not be observable in such cases.

The approach presented in this study was developed to overcome these limitations and perform an automated laminar identification of the recording sites along a cortical silicon probe. This was achieved by finding the optimal match between the VEPs recorded across the probe in a given experiment and a template VEP profile, spanning the whole cortical thickness, that was computed by merging several recording sessions, in which the ground-true depth and laminar location of the recording sites were recovered through histology. As a result, our method was able to achieve, without the need of any subjective human judgment, a cross-validated accuracy of 79 µm in recovering the cortical depth of the recording sites and a 76% accuracy in returning their laminar position.

## Materials and methods

### Animal preparation and surgery

All animal procedures were in agreement with international and institutional standards for the care and use of animals in research and were approved by the Italian Ministry of Health: project N. DGSAF 22791-A, submitted on Sep. 7, 2015 and approved on Dec. 10, 2015 (approval N. 1254/ 2015-PR). Extracellular recording data from a total of 13 naïve, Long-Evans male rats (Charles River Laboratories), 8 from V1 sessions from (Matteucci et al. 2019) reported 5 from V1 sessions of another (unpublished) dataset, were included in the analysis. Their age ranged 3-12 months and their weight ranged from 300 to 600 g. Each rat was anesthetized with an intraperitoneal (IP) injection of a solution of 0.3 mg/kg of fentanyl (Fentanest^®^, Pfizer) and 0.3 mg/kg of medetomidin (Domitor^®^, Orion Pharma). The level of anesthesia was monitored by checking the absence of tail, ear and hind paw reflexes, as well as monitoring blood oxygenation, heart and respiratory rate through a pulse oximeter (Pulsesense-VET, Nonin). A constant flow of oxygen was delivered to the rat throughout the experiment to prevent hypoxia. A constant level of anesthesia was maintained through continuous IP infusion of the same aesthetic solution used for induction, but at a lower concentration (0.1 mg/kg/h Fentanyl and 0.1 g/kg/h Medetomidin). This was done using a syringe pump (NE-500; New Era Pump Systems). Internal temperature of the animal was thermostatically kept at 37°C using a heating pad to prevent anesthesia-induced hypothermia.

After induction, the rat was secured to a stereotaxic apparatus (Narishige, SR-5R) in flat-skull orientation (i.e., with the surface of the skull parallel to the base of the stereotax) and, following a scalp incision, a craniotomy was performed over the target area in the left hemisphere (typically, a 2×2 mm window) and the dura was removed to allow the insertion of the electrode array. The coordinates used to target V1 were ∼6.5 mm posterior from bregma and ∼4.5 mm left to the sagittal suture (i.e., AP 6.5, ML 4.5). Throughout the procedure, the eyes of the animal were protected from direct light and kept hydrated by repeated application of an ophthalmic ointment (Epigel^®^, Ceva Vetem).

Once the surgery was completed, before probe insertion, the stereotax was placed on a rotating platform and the rat’s left eye was covered with black, opaque tape, while the right eye (placed at 30 cm distance from the monitor) was immobilized using a metal eye-ring anchored to the stereotax. The platform was then rotated, so as to align the right eye with the center of the stimulus display and bring the binocular portion of its visual field to cover the left side of the display. For the whole duration of the recordings, eye and cortex were periodically irrigated using saline solution in order to keep them properly hydrated.

### Electrophysiological recordings

Extracellular recordings were performed using single-shank, 32-channel silicon probes (NeuroNexus^®^) with site recording area of 775 μm^2^ and 25 μm of inter-site spacing. After grounding (by wiring the probe to the animal’s head skin), the electrode was manually lowered into the cortical tissue using an oil hydraulic micromanipulator (Narishige, MO-10; typical insertion speed: ∼ 5 μm/s), up to the chosen insertion depth (∼800-1200 μm from the cortical surface). The probes were inserted with a variable tilt, between 0° and 30°, relative to the surface of the skull. It should be noted that, in general, such tilt does not exactly correspond to the tilt of the probe insertion track with respect to the perpendicular to the pial surface at the insertion point (see Fig. 3B), because of the curvature of the brain surface. Extracellular signals were acquired using a system three workstation (Tucker-Davis Technologies) with a sampling rate of 25 kHz. Before insertion, the probe was coated with Vybrant^®^ DiI cell-labelling solution (Invitrogen, Oregon, USA) to allow visualizing the probe insertion track post-mortem through histological procedures. To this aim, at the end of the recording session, an electrolytic lesion was also performed by delivering current (5 μA for 2 seconds) through the 4 deepest channels at the tip of the shank.

Raw voltage traces were acquired at 25 kHz sampling rate and later downsampled to 610 Hz after lowpass filtering to obtain LFPs. Traces were then visually inspected to identify possibly “broken” channels (easily identifiable by having very strongly attenuated voltage variations as compared to the surrounding channels). The traces recorded at such defective sites were replaced by the average of the two surrounding channels (i.e., above and below the broken site), so as to obtain a set of LFP traces without artifactual signal discontinuities across channels.

After this pre-processing step we extracted 168 ms-long (i.e. 275 samples-long) VEP traces from each site/channel, each starting from the onset of stimulus presentation. More specifically, the responses to all the repeated presentations of all the drifting gratings used during a recording session were averaged to obtain a smooth VEP for each channel (see next section and the Results).

### Visual stimuli

During a recording session, two kinds of visual stimulation protocols were administered to the rat.

Initially, a 15 min-long receptive field (RF) mapping procedure was used to verify in real-time the identity of the targeted area (based on the assessment of the known retinotopy of rat V1) and to optimize the location of the RF centers for the following, main stimulation protocol (i.e., to ensure that most RFs fell inside the monitor, by rotating the platform or repositioning the eye through adjustments of the eye-ring). Such a brief RF mapping protocol and its use for visual areas identification has been thoroughly described elsewhere.

Once the probe was positioned at the final recording location, the main presentation protocol was administered. For the 8 animals taken from (Matteucci et al. 2019), this included 1 s-long drifting gratings, made of all possible combinations of 3 spatial frequencies (SF; 0.02, 0.04 and 0.08 cycle/°), 3 temporal frequencies (TF; 2, 4 and 8 Hz), and 12 directions (from 0° to 330°, in 30° increments). For the 5 additional animals from the other dataset, the main protocol included 1 s-long drifting gratings, made of all possible combinations of 2 spatial frequencies (SF; 0.02, 0.04 cycle/°), 2 temporal frequencies (TF; 2, 6 Hz), and 12 directions (from 0° to 330°, in 30° increments). Each grating stimulus was presented in 20 repeated trials. All stimulus conditions were randomly interleaved, with a 1 s-long inter stimulus interval (ISI), during which the display was set to a uniform, middle-gray luminance level.

Stimuli were generated and controlled in MATLAB^®^ using the Psychophysics Toolbox package, and displayed with gamma correction on a 47-inch LCD monitor (SHARP PNE471R) with 1920×1080 pixel resolution, 220 cd/m^2^ maximum brightness and spanning a visual angle of 110° azimuth and 60° elevation. Grating stimuli were presented at 60 Hz refresh rate.

### Histology

At the end of each recording session, the animal was deeply anesthetized with an overdose of urethane (1.5 gr/kg) and perfused transcardially with phosphate buffer saline (PBS) 0.1 M, followed by 4% paraformaldehyde (PFA) in PBS 0.1 M, pH 7.2. The brain was then removed from the skull, post-fixed in 4% PFA for 24 h at 4°C, and then immersed in cryoprotectant solution (30% w/v sucrose in PBS 0.1 M) for at least 48 h at 4 °C. The brain was finally sectioned into 30 μm-thick coronal slices using a freezing microtome (Leica SM2000R, Nussloch, Germany). Sections were mounted immediately on Superfrost Plus slides and let dry at room temperature overnight. A brief wash in distilled water was performed to remove the excess of crystal salt sedimented on the slices, before inspecting them at the epifluorescence microscope. Each slice was then photographed with a digital camera (MBF Bioscience CX9000) adapted to a Leica microscope (Leica DM6000B-CTR6000, Nussloch, Germany), acquiring both a DiI fluorescence image (700 nm DiI filter) and a brightfield image, using a Leica PL Fluorotar 2.5X/0.07 objective. Following the acquisition of this set of images, the sections displaying the electrode fluorescent track were further stained for Nissl substance using a 0.5% Cresyl Violet Acetate solution, and new pictures were taken at 2.5X magnification. By superimposing the fluorescence, bright-field and Nissl-stained images, it was possible to reconstruct the tilt and the anteroposterior (AP) position of the probe during the recording session, as well as the cortical depth and laminar location of all the recording sites. Specifically, the boundaries between the cortical layers were identified, based on the difference in size, morphology and density of the Nissl-labelled cells across the cortical thickness. The position of the probe relative to such boundaries and to cortical surface was determined by tracing the outline of the fluorescent track, and taking into account, when available, the location of the electrolytic lesion performed at the end of the recording session. Based on the known geometry of the silicon probe, it was possible to infer the location of each recording site along the shank, thus estimating its cortical depth and laminar location. The former was always measured along the line perpendicular to the layers at the site of interest. Such analysis was carried out using Inkscape 0.48.3.1. For illustrative purposes (Fig. 3B), we acquired large, higher magnification images of some slices from a selected representative session. To do so, we used a motorized inverted confocal Nikon Eclipse TI microscope equipped with a digital camera (Hamamatsu C4742-95), with a 20X/ 0.5 (Nikon Plan Fluor) objective. Image acquisition and stitching of a large field of 7×7 mm with 40% overlap was handled by Nikon NIS-Elements AR 4.0 software. Images were cropped and sized using Adobe Photoshop CS6 and the montages were generated in Adobe Illustrator CS6.

## Results

Our automated method for inferring the cortical depth and laminar location of the recording sites of a silicon probe is based on three key steps. First, we had to build a “template VEP profile” along the cortical thickness, by averaging the waveforms of the VEPs recorded at the same cortical depth across multiple, repeated experimental sessions employing the same (or similar) visual stimulation protocol. Second, we had to establish a map between cortical depth and laminar location in the primary visual cortex (V1) of the animal model used to demonstrate our method – the Long-Evans rat. Third, we had to find the optimal match between the template VEP profile and candidate spatial arrangements of the VEPs recorded from a given probe in a given session, so as to infer the cortical depth of the recording sites of the probe and, through the depth-to-layer map, their laminar location.

This required obtaining a rich dataset of VEPs, coupled with the histological depth localization of the electrodes from which they were recorded. This was achieved by merging the V1 recordings collected in (Matteucci et al. 2019), for which the histological analysis yielded the most accurate estimates (i.e., 10 recording sessions performed in 8 different rats), with 8 additional V1 recording sessions obtained from 5 rats (part of a dataset collected for another study that is currently in preparation). Both datasets consisted of extracellular recordings performed in anesthetized, adult, Long Evans male rats, passively exposed to full-contrast, sinewave drifting gratings of different orientations, spatial and temporal frequencies, each lasting 1 s and presented in 20 repeated trials (see Materials and method). The electrode arrays used to perform these recordings were single-shanks Neuronexus^®^ silicon probes, with either 32 or 64 recording sites and 25 µm inter-site spacing. Raw voltage traces were acquired at 24 kHz sampling rate and later downsampled to 610 Hz after lowpass filtering, to obtain LFPs (see Materials and methods).

### Building the template VEP profile and the depth-to-layer map

To obtain the average waveforms needed to build the template VEP profile, 168 ms-long segments (corresponding to 275 samples) were extracted from the LFP traces that were recorded in response to all the presentations of the drifting gratings (starting from the onset of each stimulus). For any given recording site, these 168 ms-long VEPs were averaged across all presented trials, directions, spatial and temporal frequencies, so as to average out the large trial-by-trial variability that is typical of ERPs and let the common, depth/layer-specific dynamics emerge. It should be noted here that this averaging will also wash out features of the VEP waveforms that would possibly distinguish the responses produced by different stimuli. In our method this is intentional, since our template-matching procedure will mostly relie on the amplitude, width and polarity of the initial, very strong LFP deflection that is invariably induced by a high-contrast stimulus presented in the visual field of an animal (regardless of finer visual properties – e.g., the orientation of a grating). As a matter of fact, the highly stereotyped deflection at the onset of VEP responses has made them, throughout the years, a useful measure of the magnitude of neural mass activity – e.g., to assess electrophysiological perceptual thresholds, such as contrast thresholds or spatial acuity (Kang et al. 2013; Porciatti et al. 1999; Tokashiki et al. 2018). The reliance on such robust and largely stimulus-invariant physiological signal is one of the strengths of our method, since it allows its application to datasets collected for other purposes, without the need of designing and presenting additional, ad-hoc visual stimuli.

After obtaining the average VEP for every channel of each probe used in our experiments, we further averaged the resulting waveforms based on the depth (i.e., the distance from the cortical surface) at which they were recorded. To this aim, we discretized the cortical thickness (that, in rat V1, approximately spans 1350 µm (Swanson 2004)) into nine 150 µm-wide bins and we averaged the VEPs recorded from all the sites whose depth, as measured though histology, fell inside the same bin. This yielded the template VEP profile shown in Fig. 1A, where a small, upward deflection, starting at about 90 ms following stimulus presentation, gradually molds into an increasingly deeper, broader and later downward deflection, while traveling from the surface to the bottom of the cortex – a profile that is qualitatively consistent with the laminar pattern of VEPs reported in previous rodent studies (Deliano et al. 2018; Sieben et al. 2013).

**Figure 1.**
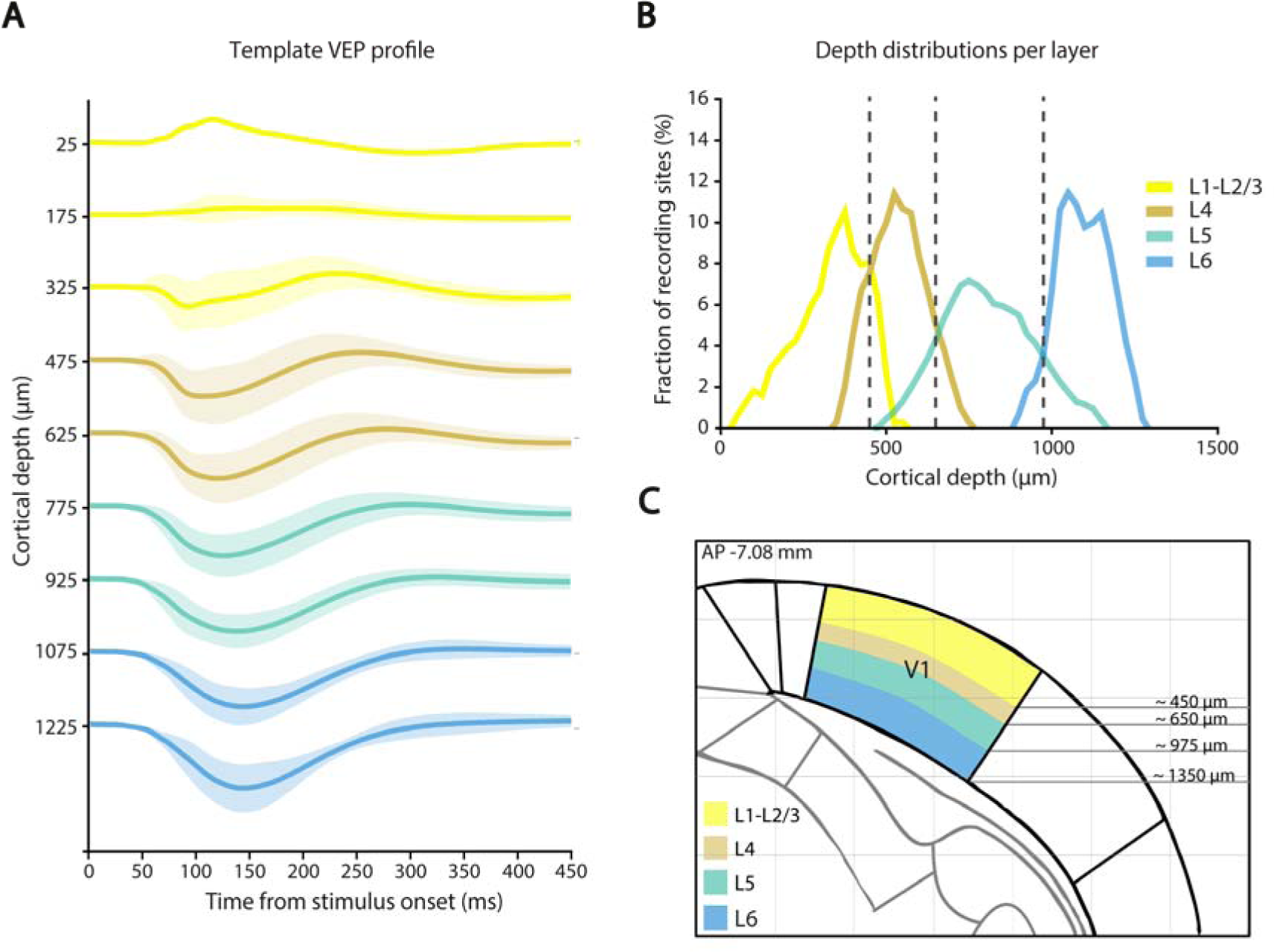
Template VEP profile and depth-to-layer map. A) The average waveforms (thick lines) ± SEM (shaded areas) of the VEPs recorded across all the 18 sessions of our experiment are plotted as a function of the cortical depth. Every waveform is the average of all the VEPs falling within a given 150 µm-wide cortical span. The colors label the cortical layers to which the waveforms belong, according to the map established in B-C (see key in B). B) Each line shows the empirical probability that a recording site within a given cortical layer was located at a given cortical depth (with both the depth and layer attribution recovered from the Nissl-stained brain slices obtained for the 18 recording sessions; see example in Fig. 3C). The dashed lines indicate the optimal boundaries between pairs of adjacent layers, as defined by taking the depths at which the corresponding distributions intersected. C) The span of each cortical layer or group of cortical layers (as defined in B) is color coded (see key) and superimposed to the outline of a coronal section of the rat brain derived from the Paxinos and Watson atlas (Paxinos and Watson 2013). AP stands for anteroposterior, and refers to the distance of the coronal section from bregma in mm.

To obtain the depth-to-layer map, we established the laminar location of the recording sites in each recording session through visual inspection of the corresponding Nissl-stained brain slice (see Materials and methods for details and Fig. 3B for an example slice). By combining this information with the depth of the sites (also recovered from the Nissl sections), we computed the probability, for a site within a given cortical lamina, to be found at a given cortical depth. Fig. 1B shows the resulting depth distributions for layers 1-3 (yellow), layer 4 (brown), layer 5 (green) and layer 6 (blue). Given these distributions, the optimal boundary between a pair of adjacent layers can be defined as the depth at which the corresponding distributions intersect, since this choice minimizes the number of incorrect attributions between the two layers. The resulting boundaries (dashed lines) were thus used to build the final depth-to-layer map, shown in Fig. 1C.

### The VEP template-matching algorithm

At the heart of our depth inference method there is a template-matching algorithm, whose aim is to infer the most likely insertion depth and tilt of each shank of the silicon probe used in a given recording session, relative to the surface of the cortex. The algorithm consists of two steps. First, we compute the Euclidean distance between the observed input data (i.e., the VEPs recorded across the channels of the shank under exam) and the VEPs expected for any possible combination of depths and tilts of the shank over a 25×25 search grid. The ranges of hypothetical depth and tilt values considered in building such a grid (i.e., a 400-1600 µm range for tip depths and a 0°-50° range for tilts) are chosen so as to roughly match those of for our electrode insertions. The expected VEPs are obtained from the template VEP profile (Fig. 1A) under the hypotheses that the tip of the shank (i.e., the deepest recording site) is positioned at the selected depth and the whole shank is rotated of the selected tilt with respect to the cortical surface. These tilt and depth parameters, combined with the known inter-site spacing, univocally specify the expected depth of each recording site and, therefore, the expected VEP associated to that site, based on the template VEP profile. Fig. 2 graphically illustrates this procedure, by showing how two hypothetical shank insertions, having different tip’s depths and tilts, give rise to two different patterns of expected VEPs. These are matched to the pattern of VEPs that was actually observed along the shank. The outcome of this procedure is a 25×25 matrix (Fig. 2C), where each element reports how good the match is between the observed VEPs and the expected VEPs (in terms of Euclidean distance), depending on the hypothesized insertion depth and tilt of the shank.

**Figure 2.**
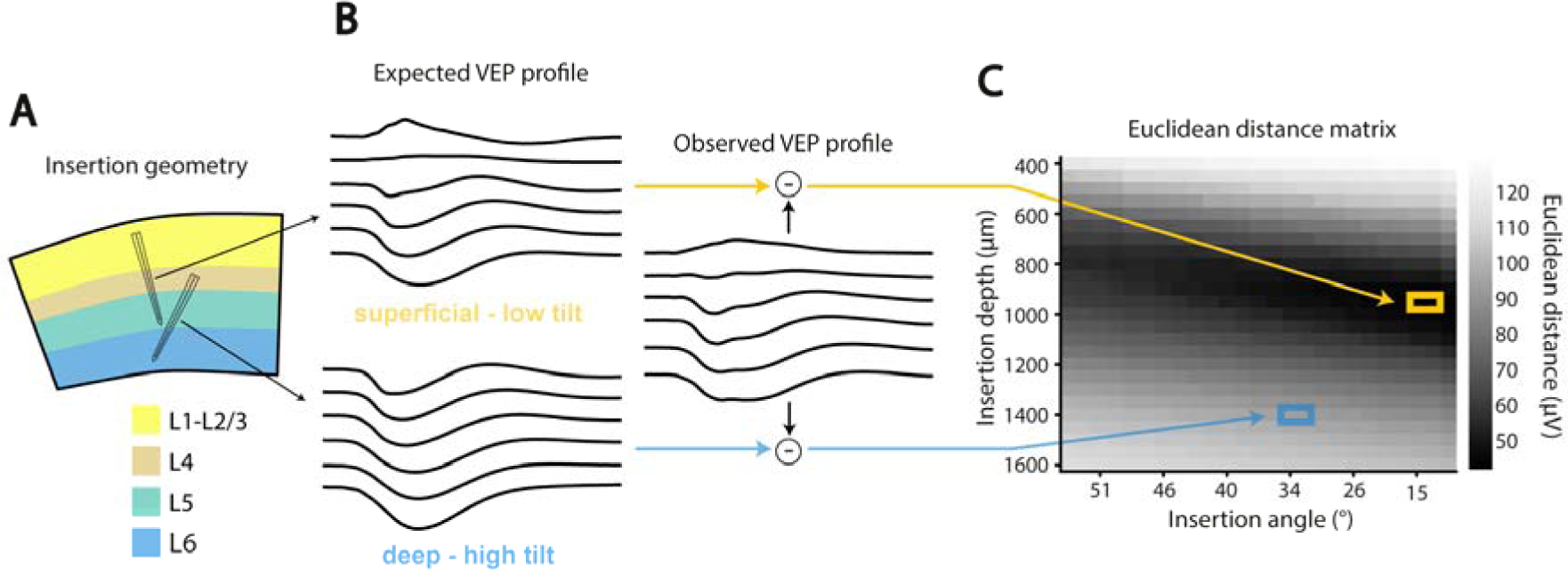
Graphical illustration of the VEP template-matching algorithm. A) The cartoon shows two hypothetical insertions of a single-shank silicon probe, reaching two different cortical depths with two different insertion angles. The cortical layers, which are color coded (see key), are the same as those shown in the depth-to-layer map of Fig. 1C. B) The hypothetical probe insertions shown in a give rise to two different patterns of expected VEPs (left), which are obtained by sampling the waveforms of the template VEP profile (shown in Fig. 1A) according to the expected depths of the recording sites. Each of the expected VEP profile is compared to the pattern of VEPs that was actually measured during the recording session (right), by computing the Euclidean distance. C) The matrix of Euclidean distances obtained by systematically considering all possible combinations of 25 insertion depths (referred to the tip of the probe and ranging from 400 to 1600 µm) and 25 insertion angles (ranging from 0 to 50°). The magnitude of the distance (in µV) is color coded (see color bar). The distances corresponding to the two insertions hypothesized in A-B are highlighted by the yellow and blue frames.

The second step of the algorithm makes use of this matrix to define the inferred insertion depth and tilt as a weighted average of all hypothesized combinations of depths and tilts, where the weights are proportional to the inverse of the Euclidean distances reported in the matrix. The resulting estimates of the insertion depth and tilt of the shank are then combined with its known inter-site spacing to predict the depth of each recording site and, using the depth-to-layer map shown in Fig. 1C, its layer assignment. As shown in the example of Fig. 2C, a quite broad “valley” (dark region) of small Euclidean distance values was typically observed across the depth/tilt plane, indicating the compatibility of the recorded VEP profile with an extended range of plausible depth and tilt values. It was because of the broadness of the compatibility region that the inferred insertion parameters were computed as a weighted average over all possible depth and tilt values, instead of simply taking the minimum over the Euclidean distance matrix. This prevented relying disproportionately on the contribution of a single combination of insertion parameters, discarding other, almost equally likely candidates (i.e., those yielding only slightly higher Euclidean distance values). We verified a posteriori that the weighted averaging was indeed superior to selecting the minimum of the Euclidean distance matrix, since the former approach yielded estimates of the recording sites’ depth (shown in Fig. 5A; see the following sections) that were twice as accurate as those obtained with the latter method.

### Validation of the depth inference method

To measure the accuracy of our method at inferring the cortical depth of the recording sites and their laminar location, we used a leave-one-out cross-validation procedure that worked as follows. We took only 17, out of the 18 recording sessions, to build the template VEP profile (see Fig. 1A), and we used it to predict the depth of the recording sites for the remaining left-out session. This procedure was applied exhaustively, so as to obtain the cross-validated accuracy of our predictions for each of the 18 sessions. This approach, which is widely used in the field of machine learning, is extremely data efficient, since it allows splitting the data into a training and a test sets, thus testing the capability of the method to generalize to new data, while maximizing the number of sessions used to build the template in each particular split (i.e., all but one sessions).

Fig. 3 shows the reconstruction of the recording sites of a 32-channel silicon probe, obtained with our template-matching method for an example session. This session was chosen for illustrative purposes, because the error in inferring the depth of the sites was the largest among the 18 sessions and, as such, it allowed a clearer visualization of the difference between prediction and ground-truth. Yet, for each of the 32 sites, the predicted depth (dots) was at less than 212 µm from the actual depth (diamonds), with an overall root mean squared error (RMSE) of 166 µm (the prediction was based on the shank insertion geometry that was inferred from the matrix of Euclidean distances shown in Fig. 2C). The actual depth was established by visual inspection of the Nissl-stained histological section (Fig. 3B), with superimposed the fluorescence image of the insertion track of the probe (in red), which had been coated with the fluorescent dye DiI before starting the recording (see Materials and methods). The consistency between the mechanical lesion left in the tissue by probe insertion (as inspected in Nissl stained sections) and the DiI staining enabled robust visual identification of probe track. Indeed, thanks to is capability to allow precise labeling of the tissue around the probe with very little diffusion (as also visible in Fig. 3B), the use of fluorescent dyes like DiI has become a standard approach for marking microelectrode penetration tracks in acute recording experiments (Blanche et al. 2005; DiCarlo et al. 1996; Niell and Stryker 2008; Tafazoli et al. 2017). In addition, in some of the sessions (about 1/3), electrolytic lesions, produced by delivering current through multiple channels of the probe at the end of the recording session, further eased the accurate identification of the probe insertion track (see Materials and methods).

**Figure 3.**
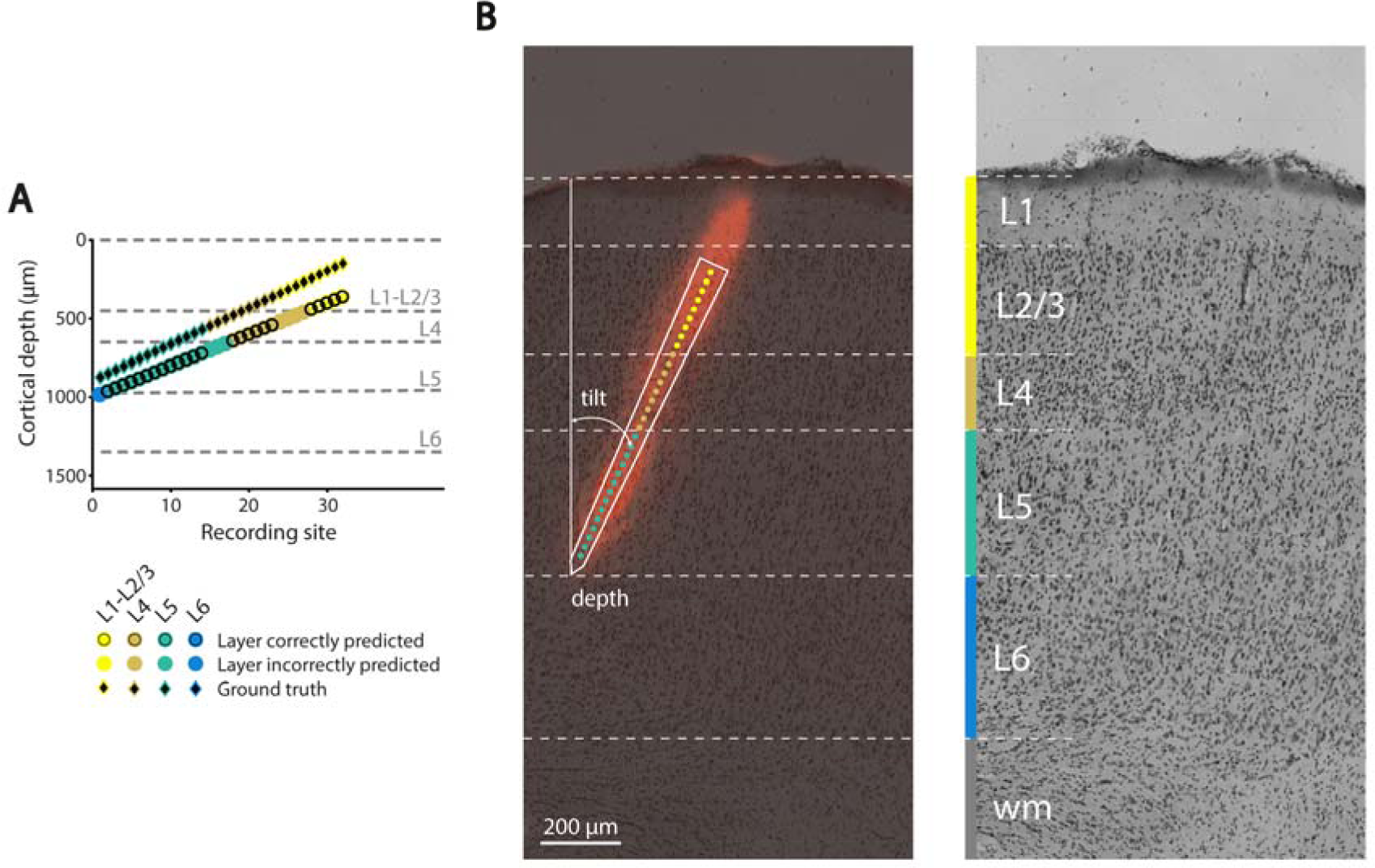
Validation of the VEP template-matching algorithm. (A) The depth of the recording sites of a single-shank silicon probe, as predicted by our VEP template-matching algorithm (colored dots), is compared to the actual depth of the sites, as recovered through histology (black diamonds). The dashed lines show the boundaries of the cortical layers, as established by the depth-to-layer map of Fig. 1C, and the color of the dots indicates the predicted layer of each recording site (see key), according to such map. The circular, black frame around a dot indicates that the layer prediction was correct, when compared to the actual laminar location of the recording site (color coded by the frame placed around each of the black diamond) that was obtained by histological analysis of the corresponding Nissl-stained brain slice. (B) Histological analysis of the brain slice showing the insertion track of the silicon probe analysed in A. On the left, a bright-field image of the Nissl-stained coronal slice is superimposed with a fluorescence image showing the staining (red) produced by the insertion of the probe, which was coated with the fluorescent dye DiI. From this image, it was possible to estimate the depth and insertion angle (or tilt) of the probe during the session, and, given the known geometry of the probe (white outline), it was possible to recover the depth of each recording site (colored dots). It was also possible to estimate the boundaries between the cortical layers, by inspecting an adjacent, 60 µm-apart Nissl-stained slice that did not bear the mechanical lesion produced by the insertion of the probe (image on the right). The resulting layer boundaries, which were drawn based on the variation of size, morphology and density of the Nissl-labelled cells across the cortical thickness, are marked by the white, dashed lines. These boundaries allowed establishing the laminar location of the sites that is color-coded by the frames around the black diamonds in A. WM stays for white matter. The data reported in this figure refer to the same recording session used to illustrate the VEP template-matching algorithm in Fig. 2.

Visual examination of the Nissl stained section also allowed recovering the boundaries between the cortical layers (white dashed lines in Fig. 3B), by carefully inspecting the landmark cytoarchitectonic features defining the layers themselves (i.e., packing density, size and shape of the cell bodies across the cortical depth). When such ground-truth layer attribution was compared to that predicted on the basis of the inferred cortical depth and the average depth-to-layer map of Fig. 1C (also reported in Fig. 3A – see the gray dashed lines), the fraction of sites whose laminar location (color coded in Fig. 3A) was correctly predicted (black-circled dots) was 75%.

Overall, this example session allows appreciating how two sources of errors concur to limit the accuracy of the predicted laminar location of the sites: 1) the error on the predicted depth (i.e., the vertical distance between dots and diamonds in Fig. 3A); and 2) the difference between the actual layer boundaries for a specific recording session (as assessed through histology) and those inferred from the depth-to-layer map (i.e., the difference between the white dashed lines of Fig. 3B and the gray dashed lines of Fig. 3A). Nevertheless, despite these potential error sources, we were able to achieve accurate depth and layer predictions for most of the recording sessions. This is illustrated in Fig. 4, which shows the cross-validated accuracy of our method for each of the 18 sessions examined in our study (the gray area highlights the example session previously analyzed in Fig. 3). In most cases, the absolute distance between predicted (colored dots) and measured (black dots) depth of the recording sites was lower than 115 µm (75% quantile of the absolute error distribution), with the method yielding, for some sessions, RMSEs as low as a few tens of µm. As a result, the overall distribution of absolute depth errors (across all the sites of all the recording probes) displayed a prominent peak in the 0-50 µm range, with no errors above 220 µm (Fig. 5A). This yielded a mean RSME ± SEM across sessions of 79 ± 11 µm.

Fig. 4 also reports the predicted layer attribution of the recording sites (color coded, with the layer boundaries marked by dashed lines) and whether such prediction was correct (black-circled dots), according to the histological assessment of the cortical sections. The fraction of correctly labeled sites (i.e., the accuracy of the labeling) ranged from 41% to 94%, with a mean ± SEM of 76 ± 3% across the sessions (first bar in Fig. 5B), when a distinction in four cortical laminae was considered (i.e., same as shown in Fig. 1C). When a coarser grouping of the layers in supragranular (1-3), granular (4) and infragranular (5-6) was considered, the accuracy of the labeling increased to 83 ± 3% (second bar in Fig. 5B), and further grew to 91 ± 2% for a binary partition of the layers into superficial (1-4) and deep (5-6; third bar in Fig. 5B).

**Figure 4.**
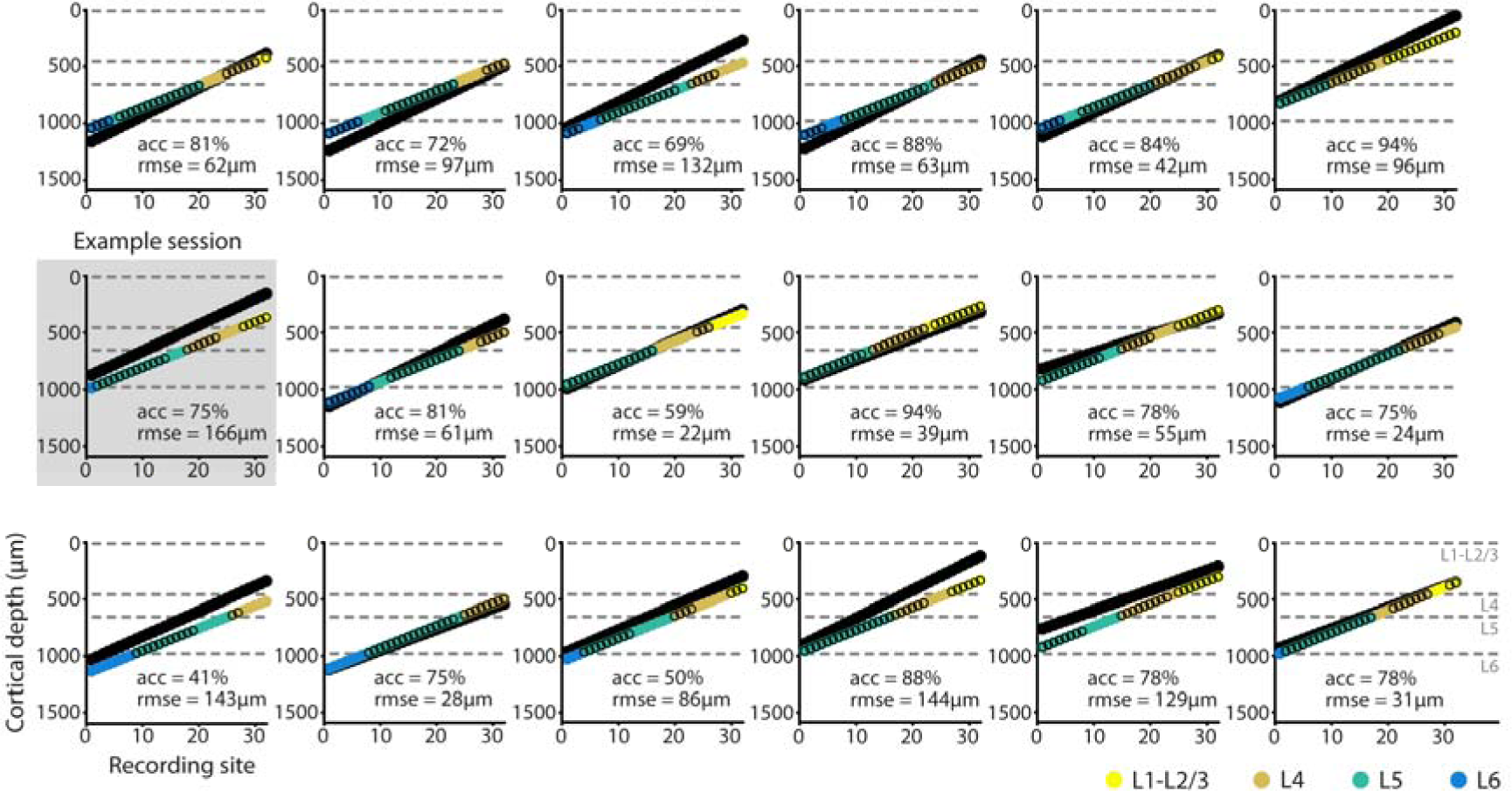
Accuracy attained by the VEP template-matching algorithm in each of the recording sessions. Every panel shows the predicted (colored dots) and measured (black dots) depths of the recording sites in a given session, along with the predicted laminar location (color-coded; same convention as in Fig. 1-3). As in Fig. 3A, a black frame around a dot indicates that the layer prediction was correct. The gray, dashed lines show the boundaries of the cortical layers, as established by the depth-to-layer map of Fig. 1C. For each session, is also reported the RMSE in inferring the depth of the sites and the accuracy in recovering their laminar location (as the percentage of correctly labelled sites). The gray area highlights the example session previously analysed in Fig. 2-3.

The errors in predicting the laminar locations were fairly homogeneously distributed across the cortical thickness, when the accuracy was measured in terms of recall (i.e., the fraction of sites belonging to a given layer that were correctly labeled as such). As shown in Fig. 5C (black bars), recall accuracy peaked in layer 5 (∼80% correct), slightly dropped in the adjacent layers (∼75%), to become ∼60% in the supragranular ones. When measured in terms of precision (i.e., the fraction of sites labeled in a given way, for which the labeling was correct), the accuracy also peaked in layer 5 (∼80% correct) and dropped substantially only in layer 6 (∼40%), likely due to the small number of recording sessions in which the probe reached this layer.

**Figure 5.**
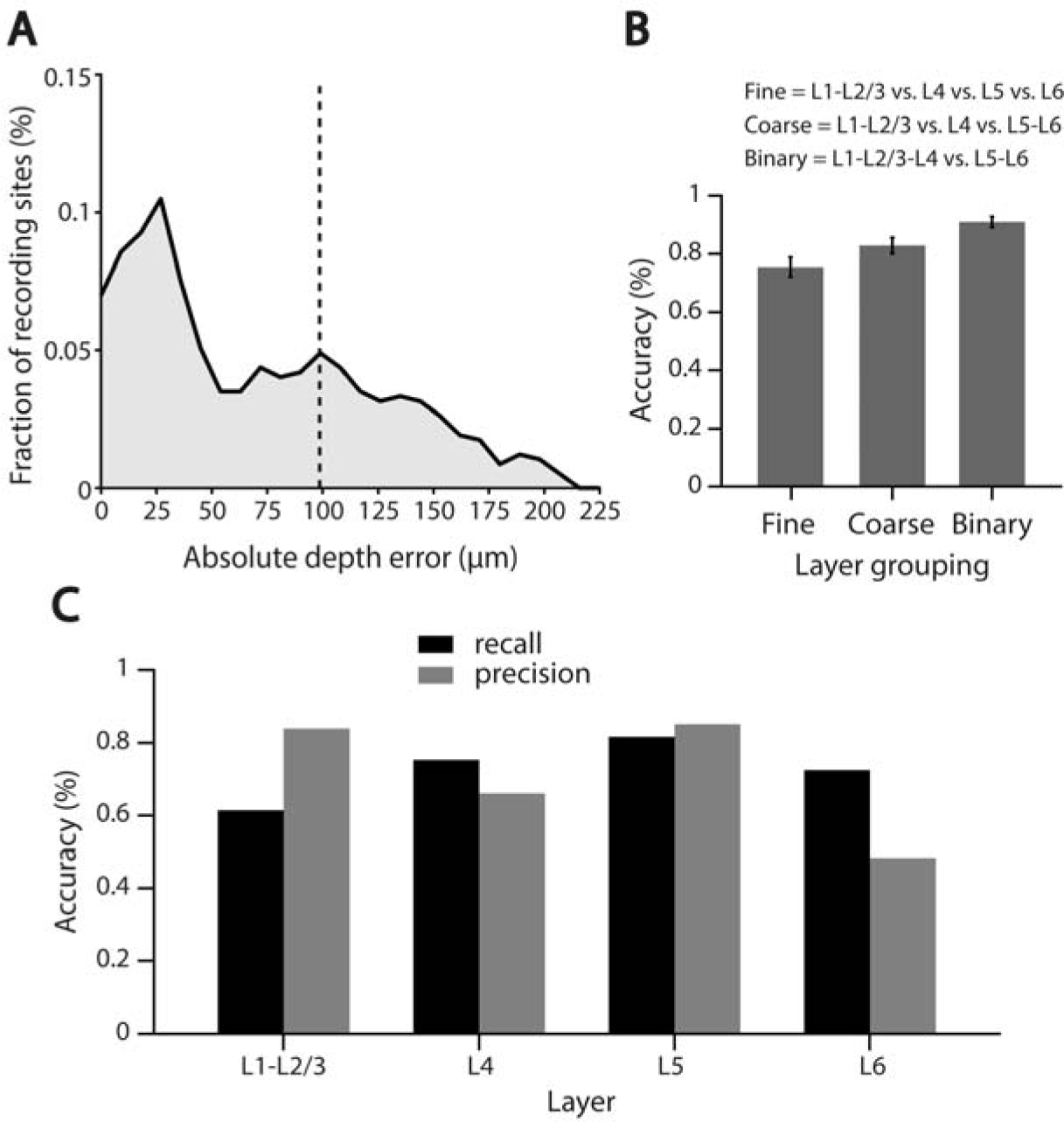
Overall accuracy of the VEP template-matching algorithm. A) Distribution of absolute errors in estimating the depth of the recording sites, as obtained by computing the absolute distance between predicted and measured depth for every site on every probe (the dashed line shows the mean of the distribution). B) Average accuracy (across the 18 recording sessions) in predicting the laminar location of the recording sites ± SEM, as measured for three different levels of distinction between the cortical layers – from fine to binary (see key). C) The overall recall (black) and precision (gray) accuracies in predicting the laminar location of the recording sites are reported as a function of the measured (for the recall) or predicted (for the precision) cortical layer of the sites. See the main text for a definition of recall and precision.

## Discussion

The method we developed provides a valuable, automated alternative for inferring cortical depth and laminar location of the recording sites along a silicon probe, as compared to approaches based on visual inspection of the pattern of current sinks and sources that can be derived from the LFPs recorded along the probe, as in CSD analysis (Dadarlat and Stryker 2017; Deliano et al. 2018; Hansen et al. 2011; Happel et al. 2010; Headley and Weinberger 2015; Hoy and Niell 2015; Li et al. 2018; Niell and Stryker 2008; Schroeder et al. 1998, 2001; Sieben et al. 2013; Takeuchi et al. 2011). One major advantage over these approaches is that, once the template VEP profile (Fig. 1A) and the depth-to-layer map (Fig. 1C) have been established from a number of training sessions (with ground-truth depth and laminar information recovered through histology), our method is fully automatized and does not require any subjective decision about the location of layer boundaries to be taken.

A second, key feature of our method is that its accuracy can be quantitatively and rigorously estimated, given the above-mentioned ground-truth information gathered through histology. Specifically, our cross-validated measurements indicate that our method is very accurate in recovering the depth of the recording sites, with an average RMSE lower than 100 µm (Fig. 5A, dashed vertical line). In addition, the method yields reliable estimates of the cortical laminae, with a 76% accuracy for fine-grained discriminations that increases to 80-90% for coarser groupings of the layers (Fig. 5B), and with both the recall and precision accuracies peaking in layer 5 (∼80% correct labeling; Fig. 5C).

It should be noted that the ground-truth laminar information used to validate our method, albeit derived from manual annotation of histological sections, is grounded on the landmark cytoarchitectonic features that define cortical layers. For this reason, laminar boundaries derived from histology are arguably the most reliable “ground truth” for the development of any algorithm aimed at predicting the laminar position of recording sites from LFPs. In other words, the histological annotation procedure is more reliable than the layer assignment resulting from either visual inspection or automated analysis of the LFP/CSD patterns recorded across the cortical depth, being the latter only an indirect proxy for the underlying anatomical structures that generate them.

It should also be noted that, unfortunately, a quantitative comparison with CSD-based methods in terms of accuracy is hard to carry out because, to the best of our knowledge, the accuracy of these approaches, relative to a ground-truth established through histology, is typically not reported. This is because inspection of the CSD patterns is usually performed in a qualitative rather than quantitative way in the neurophysiological literature. In principle, it would be possible to develop an algorithm (similar to the one described in our study) that automatically parses layer boundaries based on CSD patterns. However, being CSD derived from LFPs, it is unlikely that this method would outperform our approach, and developing such an alternative, CSD-based automated procedure was beyond the scope of our study, especially given the high accuracy we were able to achieve based on the analysis of the LFP patterns.

Another distinctive feature of our method is that it is able to reliably infer the depth and laminar location of the recording sites also in sessions where the probe did not span the whole cortical thickness and, in particular, in sessions where the VEPs were not recorded from the supragranular layers (see, in Fig. 4, the accuracy attained in sessions where the sites were all located below the boundary between layer 3 and 4). As mentioned previously, failure to sample VEPs from layers 2-3 would make it hard to properly identify the current sink associated with thalamic afferent input impinging into layer 4 by visual inspection of the CSD pattern. Obviously, also our method is sensitive to the inversion of the polarity of the VEPS in the transition from layer 3 to layer 4 – the presence of such inversion (shown in Fig. 1A) is highly beneficial, since it provides a feature with high spatial contrast in the VEP pattern observed across the cortical thickness, thus making the “valley” of Euclidean distances over the possible combinations of insertion depths and tilts deeper and narrower (as an example, see Fig. 2). Nevertheless, our method still yields a reliable estimate of the insertion parameters also when the polarity inversion is not observable, since it exploits the gradual changes in the depth, width and timing of the negative peak of the VEP trace as function of cortical depth, which are appreciable even across the deeper recoding sites (compare the brown, green and blue traces in the template of Fig. 1A).

In conclusion, we believe that our automated method for depth inference and laminar identification of recording sites represents a valuable alternative over existing, more qualitative approaches. In our study, we demonstrated the validity of our method in the primary visual cortex of the rat. Obviously, since our approach is based on obtaining a reliable estimate of the average VEP profile across the cortical thickness (Fig. 1A) and of the depth-to-layer map (Fig. 1C) through a combination of LFP recordings and histology, its application needs to be fine-tuned as a function of the cortical area, species, and brain state (e.g., awake or anesthetized) of interest. This requires collecting a fairly large dataset (e.g., 18 recording sessions in our study), thus possibly limiting the applicability of our method in cases where it is difficult and time consuming to acquire enough data – in these cases, a more qualitative approach, such as that based on visual inspection of the CSD pattern, may still be preferable. However, as already pointed out in the Results, another strength of our method is that it works with any kind of high-energy sensory stimuli that are capable of evoking strong ERPs. This means that it is not necessary to perform experiments with ad-hoc stimuli, expressly deigned to build the VEP template upon which the method rests. Rather, as we demonstrated in our study, one can “recycle” recordings, histology and stimulation protocols developed to answer different scientific questions and use such data to build the required VEP template and depth-to-layer map. This potentially extends the scope of applicability of our approach to virtually every existing dataset with a sufficiently rich pool of VEPs and ground-truth laminar locations obtained through histology. Availability of such data from previous studies would allow a research group to use them to implement our template-matching algorithm, and then use the algorithm to infer the cortical depth and laminar location of any future data collected from the same brain area/state, without the need of performing further, time-consuming histological analyses.

## Acknowledgments

We thank Rosilari Bellacosa Marotti for her help in collecting the neuronal data.

